## Supplementary material for "Two shifts in evolutionary lability underlie independent gains and losses of root-nodule symbiosis in a single clade of plants": SI Guide

**Supplementary_Notes.txt:** Supplementary Notes. A file containing supplementary notes to support the manuscript text.

**SupplementaryData1.pdf:** Supplementary phylogenetic tree. Zoomable phylogeny presented in Figure 1 with branches colored by inferred ancestral RNS status (RNS-present=blue; RNS-absent=gray).

**SupplementaryData2.pdf:** Supplementary phylogenetic trees. Twenty subsampled backbone phylogenies are presented based on ten sets of representative subsamples: Pages 1-10 were inferred using ASTRAL (page 3 is the backbone phylogeny used for subtree scaffolding and the main phylogeny presented in this paper). Pages 11-20 were inferred using RAxML-ng.

**SupplementaryData3.pdf:** Phylogenetic tree excerpted from Supplementary Data 1. Papilionoideae subtree showing enumerated gains and losses of RNS (Extended Data Tables 2 and 3). Some speciose clades are scaled, collapsed, and labeled to allow for easier viewing.

**SupplementaryData4.pdf:** Phylogenetic tree excerpted from Supplementary Data 1. Caesalpinioideae subtree showing enumerated gains and losses of RNS (Extended Data Tables 2 and 3). Some speciose clades are scaled, collapsed, and labeled to allow for easier viewing.

**SupplementaryData5.pdf:** Phylogenetic tree excerpted from Supplementary Data 1. Rosales subtree showing enumerated gains and losses of RNS (Extended Data Tables 2 and 3). Some speciose clades are scaled, collapsed, and labeled to allow for easier viewing.

**SupplementaryData6.pdf:** Phylogenetic tree excerpted from Supplementary Data 1. Fagales subtree showing enumerated gains and losses of RNS (Extended Data Tables 2 and 3). Some speciose clades are scaled, collapsed, and labeled to allow for easier viewing.

**SupplementaryData7.pdf:** Phylogenetic tree excerpted from Supplementary Data 1. Cucurbitales subtree to show enumerated gains and losses of RNS (Extended Data Tables 2 and 3). Some speciose clades are scaled, collapsed, and labeled to allow for easier viewing.

**Supplementary_Table1.xlsx**: Supplementary Table. RNS-state trait database.

**Supplementary_Table2.xlsx**: Supplementary Table. Samples used in study.
