## Supplementary Notes for "Two shifts in evolutionary lability underlie independent gains and losses of root-nodule symbiosis in a single clade of plants"

**Supplementary Note 1. Summary of NFC topology resolved in this study**

The relationships we resolved among the four NFC orders (Cucurbitales, Fabales, Fagales, and Rosales) are congruent with recent multilocus nuclear-gene based phylogenies^22^^,^^23^ that differ from earlier trees based primarily on the chloroplast genome that have often been used to analyze or discuss the evolution of SNF^6^^,^^5^^,^^24^^,^^17^. Some within-order relationships in our tree are novel but strongly supported (Supplementary Data 2, p. 3). For example, Caesalpinioideae and Papilionoideae, the two predominantly nodulating and the most speciose clades in the NFC, are typically sister (LPWG 2017), but we find instead that Caesalpinioideae is sister to the non-RNS subfamily Dialioideae. The topology used for the evolutionary models represents our current best evidence, but these relationships may be difficult to definitively resolve; for example, there was likely a near‐simultaneous evolutionary origin of all six legume subfamilies^11^.

**Supplementary Note 2. Detail on single-copy target-locus filtering**

Although we designed probes to target putatively single-copy loci, the phylogenetic breadth of the dataset, which includes ancient and recent polyploidy (e.g. Leguminosae: Wojciechowski 2019, Cannon et al. 2014; Rosaceae: Xiang et al. 2017; Cucurbitaceae: Guo et al. 2020), means that no locus is single-copy across all samples, as demonstrated by the aTRAM assembly. When multiple copies of a target gene are present in a dataset, the ideal strategy for assessing orthology would be to include all paralogs in the phylogenetic analysis. Because this was not feasible at the phylogenetic and data scales considered here, we instead designed an alternative subclade approach using phylogenetic information to reduce the likelihood of erroneously analyzing paralogs while minimizing data loss, as described below.

We performed single-copy target-locus filtering for each subclade and four outgroup samples independently (15 major clades: Fagales, Cucurbitales, seven Rosales families, one non-Leguminosae Fabales family, and five Leguminosae subfamilies; five additional clades with few species were analyzed with a closely related larger clade). For each clade, *de novo* contigs were first assembled using SPAdes (Bankevitch et al. 2012) as implemented in the target-locus assembly pipeline aTRAM2 (Allen et al. 2018). We built and used a custom aTRAM function called atram_framer.py to put the *de novo* contigs into reading frame based on the target-locus reference sequence and filtered by coverage using a 40X coverage cutoff. We then used scripts from Yang and Smith (2014) to build a phylogeny of contig sequences and identify all samples that had contigs that were not sister in the contig phylogeny; these contigs were dropped from the target-locus dataset for that clade. In cases where multiple contigs that were assembled for a single sample were sister in the phylogeny we retained the contig with the longest sequence length. Contigs that passed coverage (> 40X) and length (> 250 bp) cutoffs and were present in single copy in the contig tree were carried on to target-locus alignment and phylogenetic analysis. Scripts are available at the GitHub repository ().

**Supplementary Note 3. Detail on RNS-state database and genera that were coded as unknown in the analysis**

Nine legume genera could not be reasonably scored as RNS-present or RNS-absent because no information was available from either published observations or expert input (Supplementary Table 1) and because their phylogenetic positions in hotspots of RNS evolution did not allow for reasonably confident inference of RNS state. Species from these genera were scored as unknown in the transition rate estimation, and their states were inferred by the joint reconstruction analysis. In five cases where the inferred state would represent an independent loss or gain of RNS, we did not include these in our reported results in extended data tables 2 and 3. In Caesalpinioideae, *Arcoa* and *Tetrapterocarpon* are both inferred by ancestral character state reconstruction as RNS-present, requiring two independent gains, and *Sympetalandra* is inferred as lacking nodules, requiring an independent loss. In Papilionoideae, *Amphimas* is inferred as having nodules present, requiring an independent gain. There are unconfirmed field reports of nodulation in *Amphimas* (Diabaet et al.) that were met with skepticism by some experts (Sprent 2005), but given the present evidence, it is possible that this represents a true gain of RNS. *Petaladenium* is inferred as lacking nodules, requiring a loss.

Inferences of nodulation state for the remaining genera with unknown RNS data had no effect on inferred gains or losses: *Viguieranthus* is inferred as present but because of its phylogenetic position its inference in either state has no effect on gains or losses. However, because of its sister relationship to a genus that exhibits an independent loss, *Zapoteca*, confirmation of nodulation status in this genus is a high priority. *Diptychandra* is inferred as present, but does not require an additional gain. *Dussia* is inferred as present, as preliminarily reported by Saur et al. (2000); because of its phylogenetic position, its inference in either state has no effect on reconstruction of gains or losses.

In Caesalpinioideae, the non-monophyly of two genera, *Mora* (RNS-absent) and *Dimorphandra* (RNS-present), known previously (Legume Phylogeny Working Group 2013), precluded our genus-level scoring of RNS. To resolve this issue, the following *Dimorphandra* species were coded as lacking nodules based on phylogenetic position and no species-level observations: *Dimorphandra cuprea*, *D. ignea, D. vernicosa, D. pennigera,* and *D. polyandra*, as shown below.

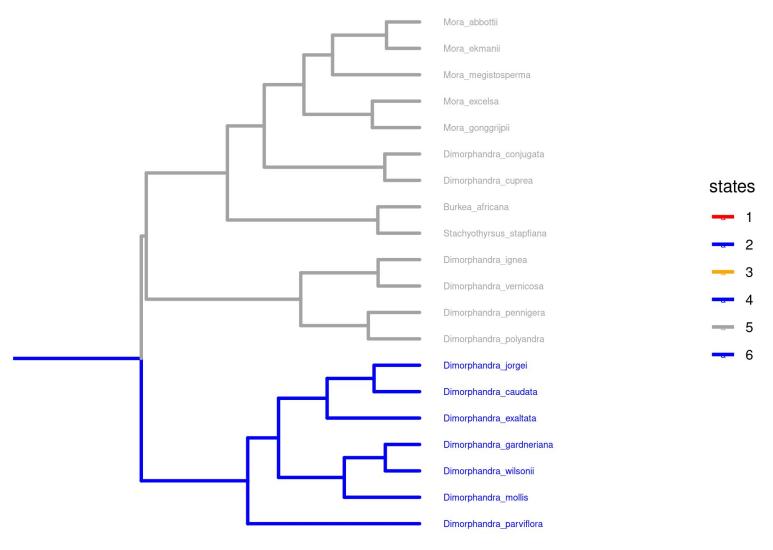

**Supplementary Note 4. Results of character state reconstruction using an alternative backbone topology of the NFC**

The relationships among the four orders of the NFC and among legume subfamilies are uncertain due to rapid divergences of these clades in evolutionary history. To check how alternative backbone topologies besides our best-evidence result affect the reconstructed history of RNS, we merged subtrees for these orders and subfamilies together onto a backbone representing relationships commonly recovered in previous studies and used to study RNS. Relationships among orders were set according to APG IV (2016), i.e., (((Fagales,Rosales),Cucurbitales),Fabales), largely representing chloroplast phylogenetic information, and relationships among legume subfamilies were set according to Koenen et al. 2020. The analysis of Koenen et al. (2020) omitted the legume subfamily Duparquetioideae and the Fabales family Surianaceae; we inserted these in positions congruent with our main phylogeny. We reran our hidden state estimation and reconstruction on this phylogeny using identical methods. The overall result (Supplementary Fig. 3A) is highly consistent with the reconstruction using the best-evidence topology recovered with our data: the ancestor of the NFC is in the deep-precursor state, and the same number of independent origins of SNF is hypothesized (Extended Data Table 2). One notable difference is that the direct-precursor state is not recovered at such a high rate (compare Figure 2 and Supplementary Fig. 3B), meaning the reconstructed state is not as transient and therefore is seen more often in the character state reconstruction.
