## Supplementary figures and images for "Two shifts in evolutionary lability underlie independent gains and losses of root-nodule symbiosis in a single clade of plants"

### Supplementary Data 1

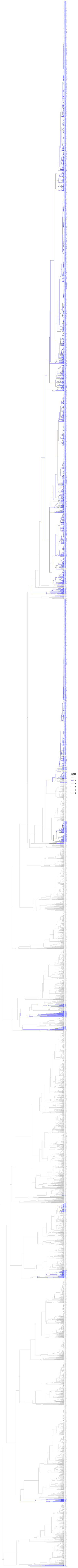

### Supplementary Data 2

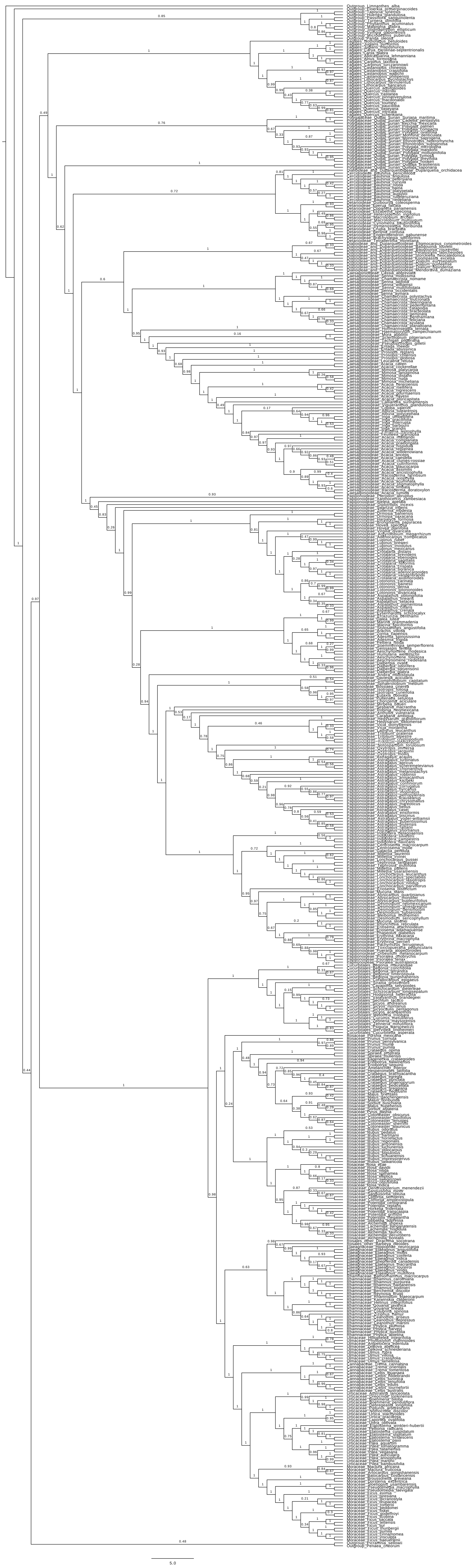

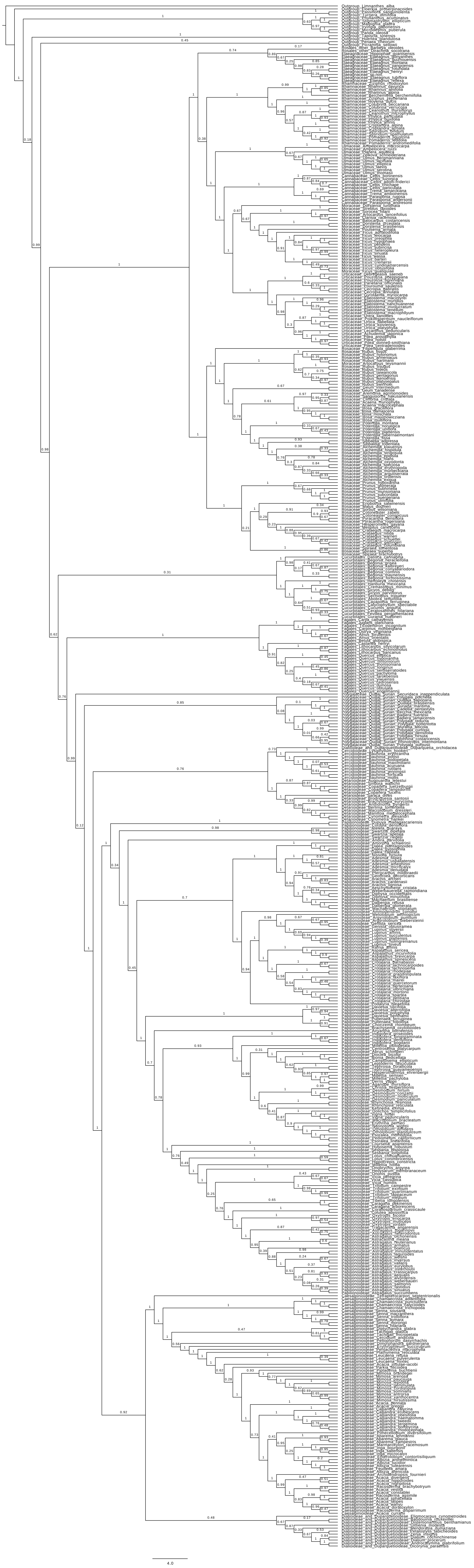

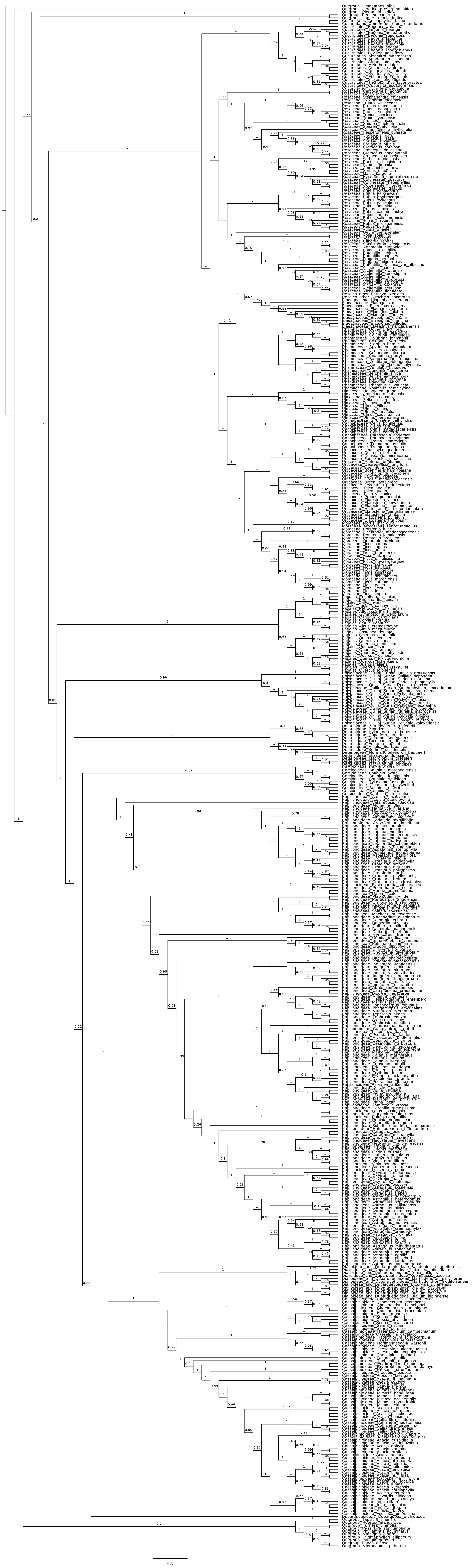

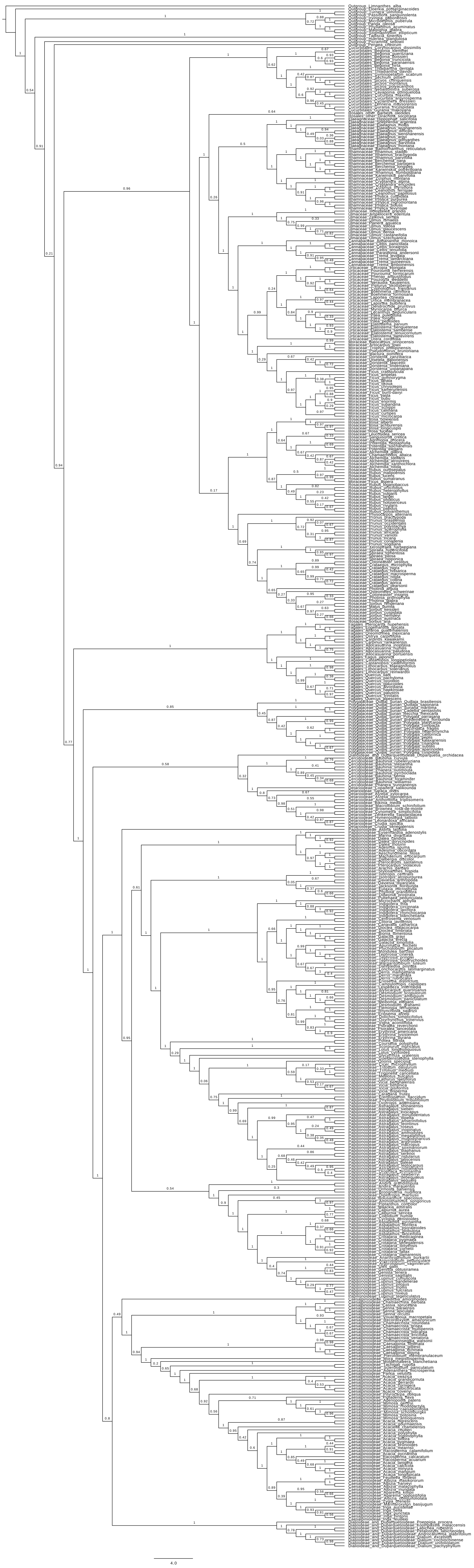

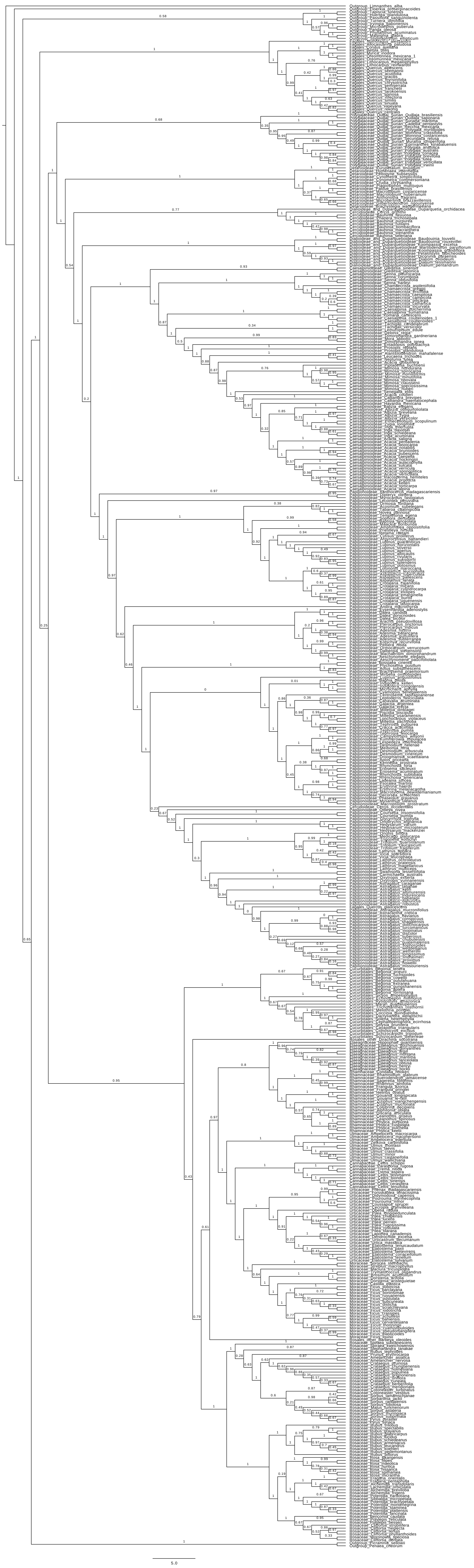

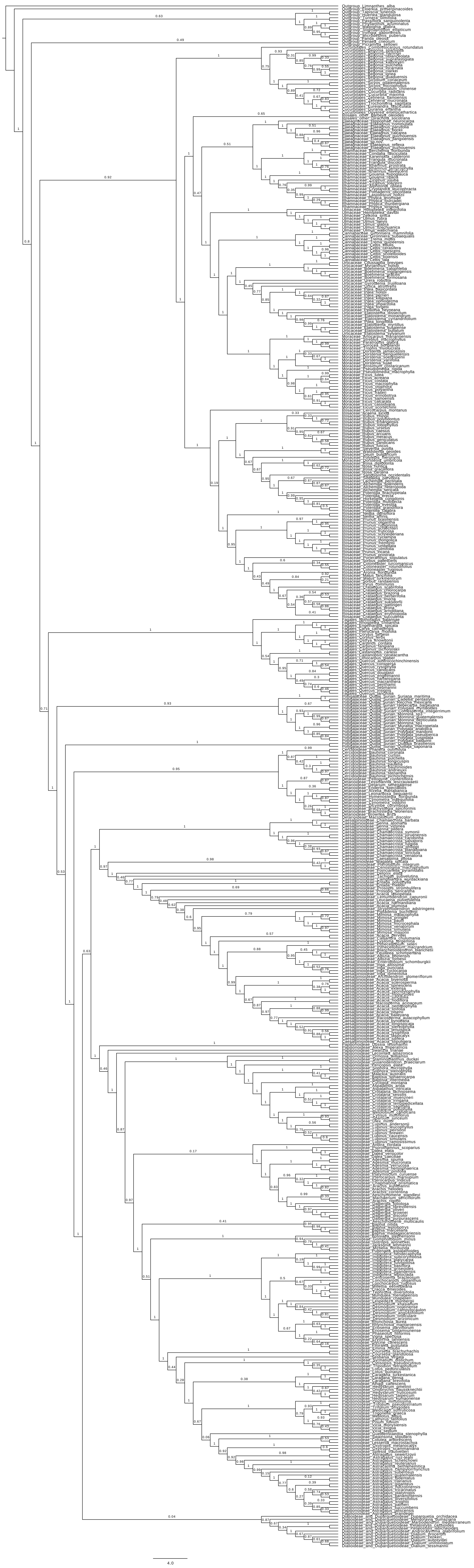

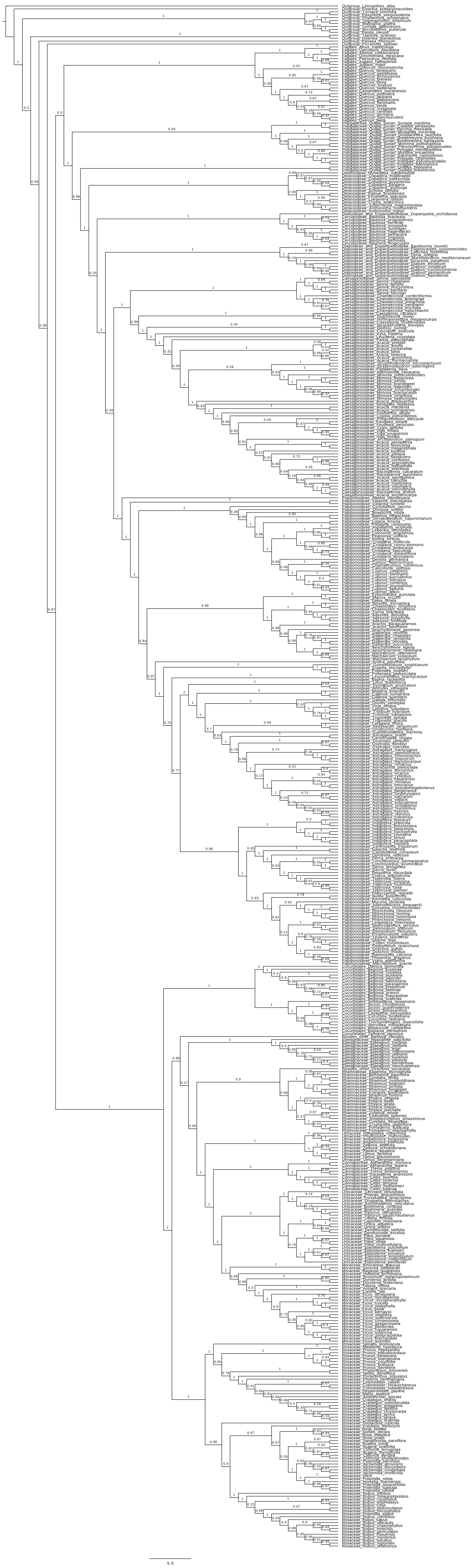

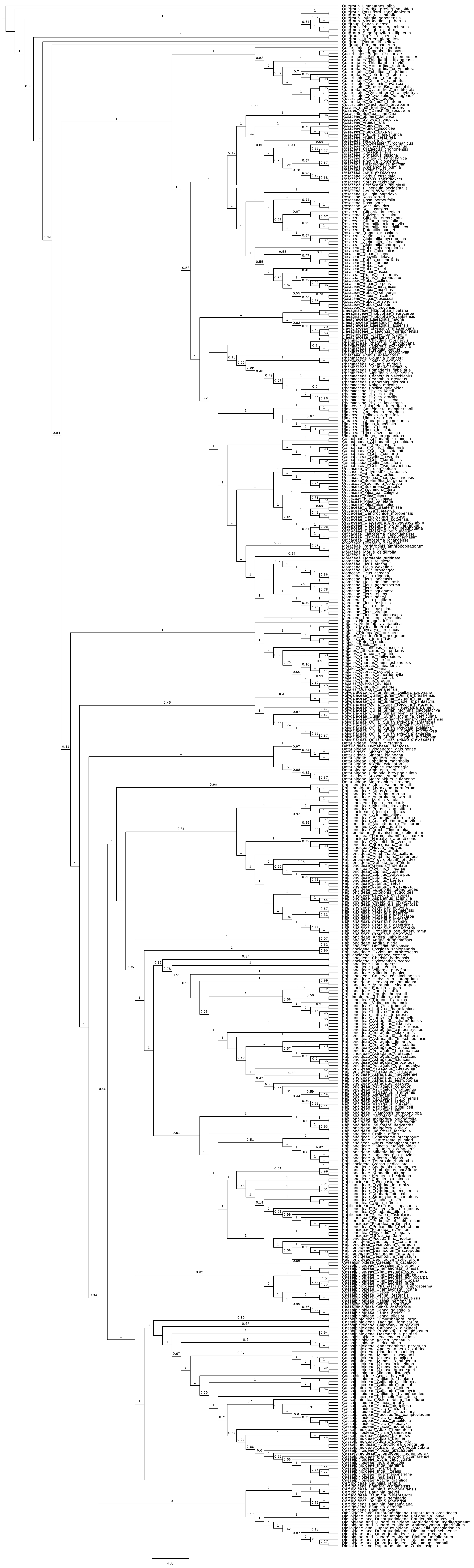

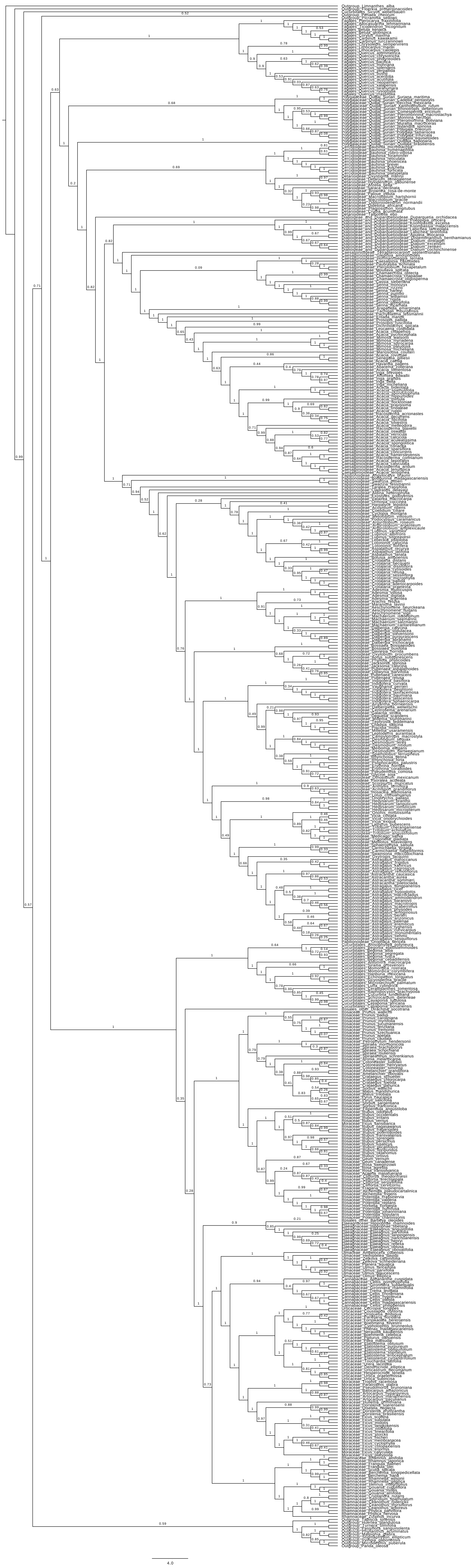

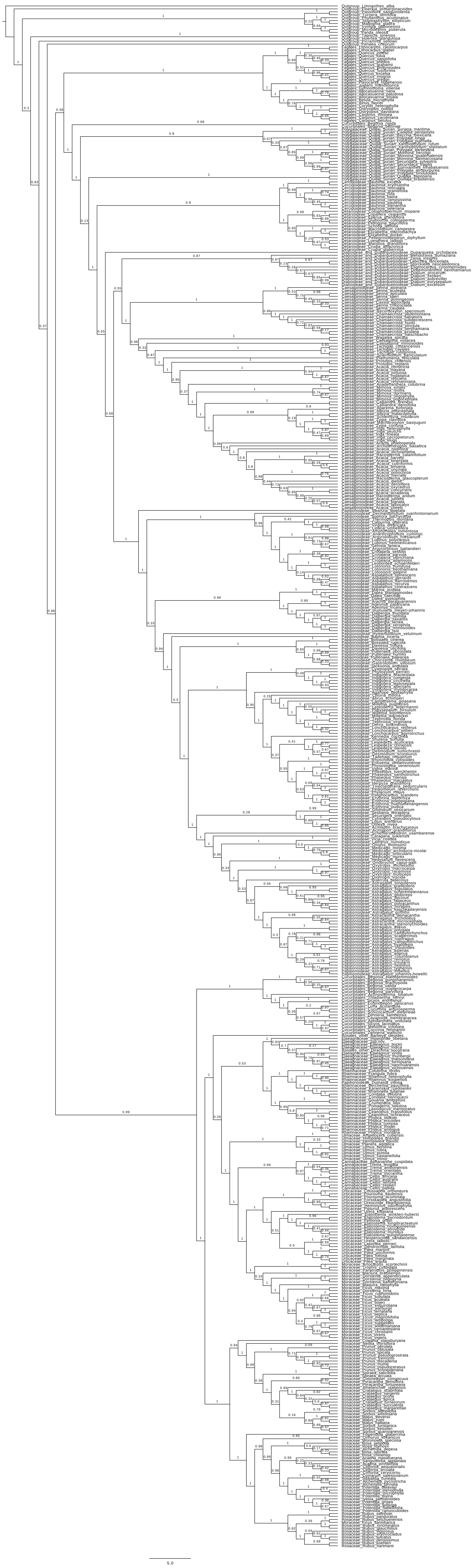

### Supplementary Data 3

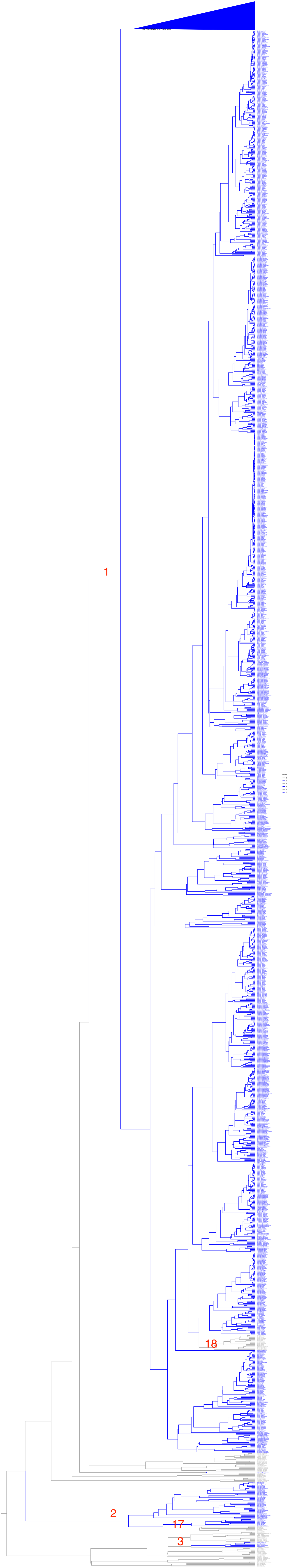

### Supplementary Data 4

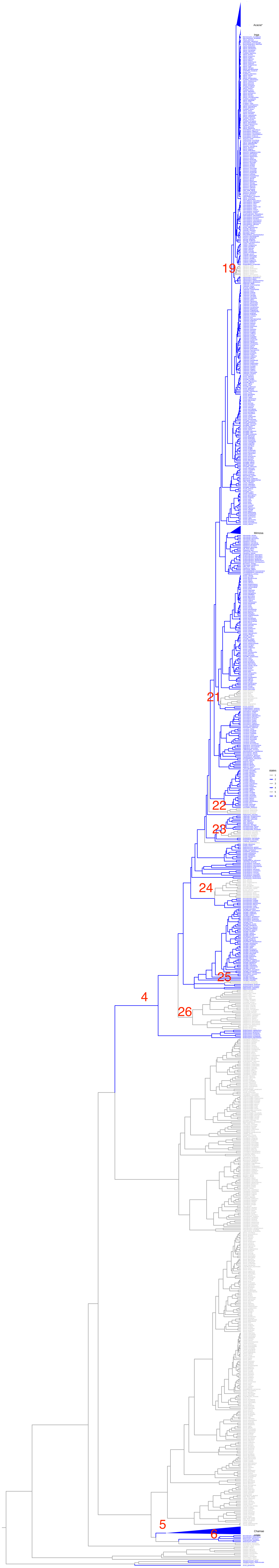

### Supplementary Data 5

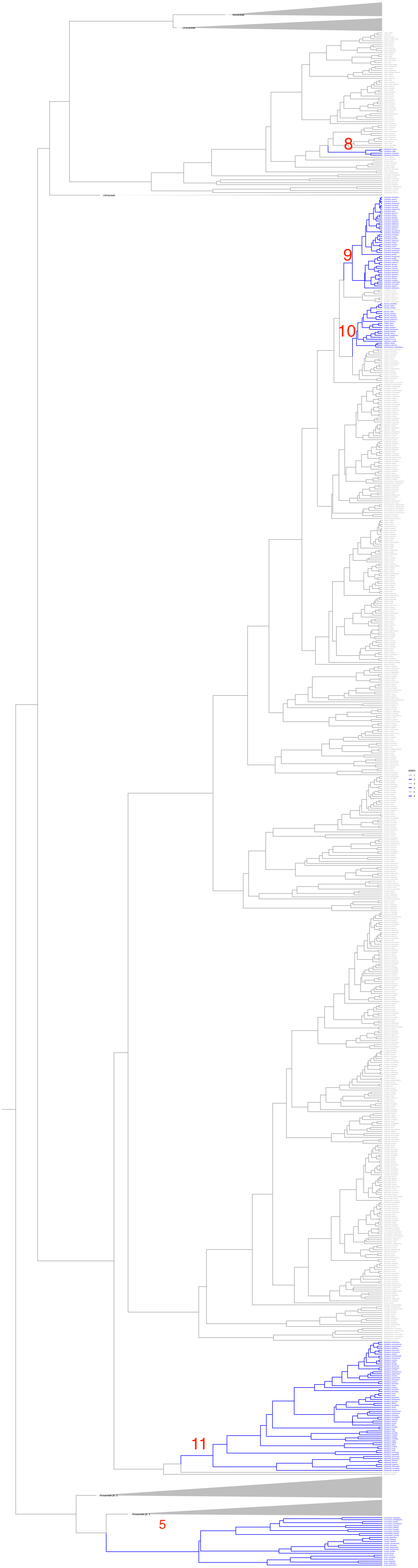

### Supplementary Data 6

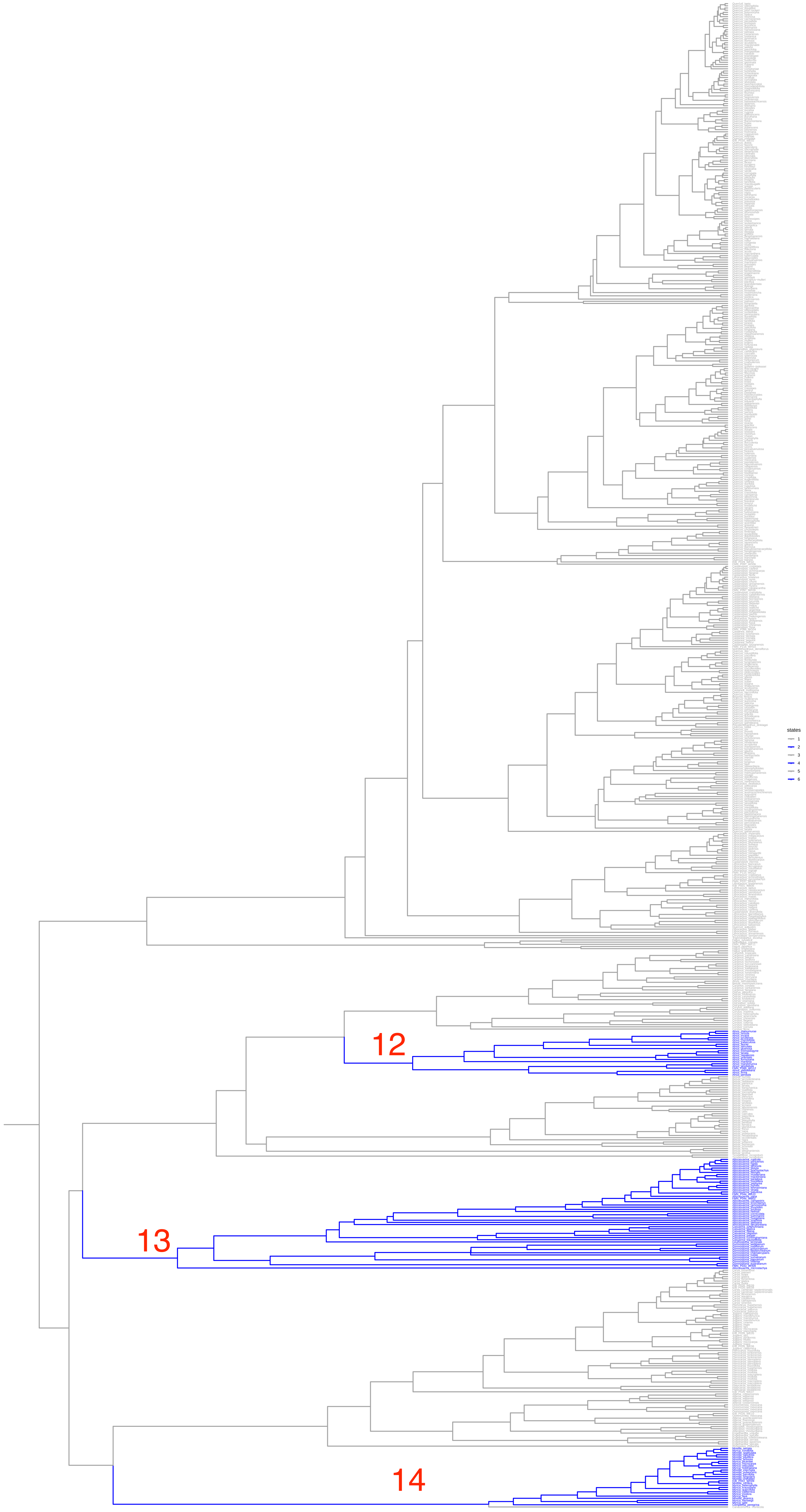

### Supplementary Data 7

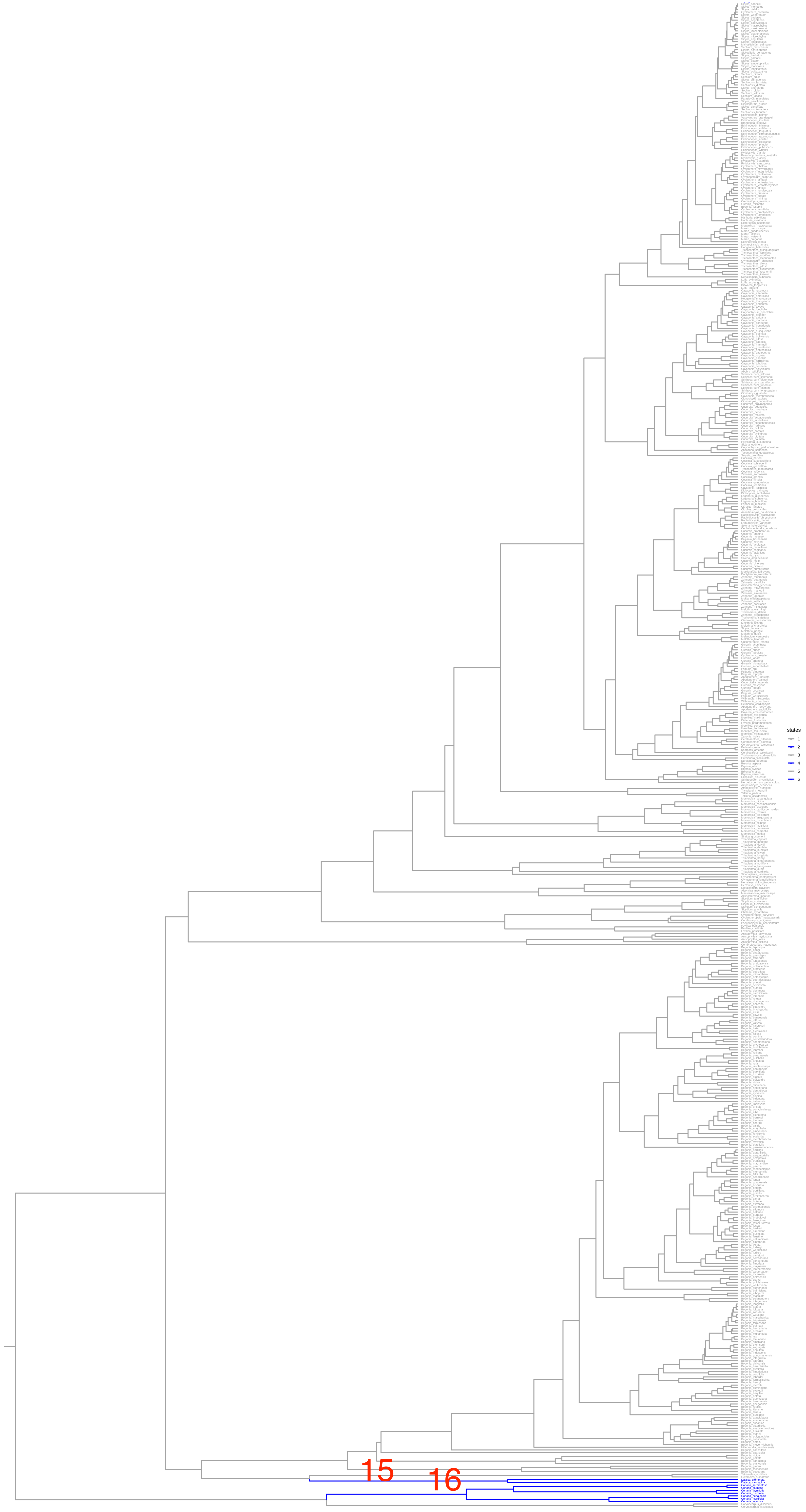
