## Extended Data Tables and Figures for "Two shifts in evolutionary lability underlie independent gains and losses of root-nodule symbiosis in a single clade of plants"

**Extended Data Table 1.** Results of model fit tests for the three analyses (A-C). The best model for each analysis is highlighted in green.

**A.** Results of model fit test for the main analysis testing one to five hidden rate categories. (The model-fit of the three-state two-rate model is reported here.)

| Number of Rate Categories | "Precursor" state? | AICc | deltaAIC |
|---|---|---|---|
| 1 | NA | 466.491 | 53.61 |
| 2 | No | 415.33 | 2.443 |
| 2 | Yes | 413.95 | 1.063 |
| 2 ; 3 state | Yes | 462.063 | 49.176 |
| 3 | No | 415.074 | 2.187 |
| 3 | Yes | 412.887 | 0 |
| 4 | No | 426.421 | 13.534 |
| 4 | Yes | 451.759 | 38.872 |
| 5 | No | 433.859 | 20.972 |
| 5 | Yes | 453.143 | 40.256 |

**B.** Results of model fit tests for fixed-ancestral-state analysis.

| Number of Rate Categories | NFC Ancestor | "Precursor" state? | logLik | AICc | deltaAIC |
|---|---|---|---|---|---|
| 1 | RNS-present | NA | -249.007 | 502.014 | 83.244 |
| 2 | RNS-present | No | -209.77 | 431.555 | 12.785 |
| 2 | RNS-present | Yes | -211.37 | 432.751 | 13.981 |
| 3 | RNS-present | No | -202.302 | 428.605 | 9.835 |
| 3 | RNS-present | Yes | -211.867 | 418.77 | 0 |
| 3 | RNS-absent | Yes | -195.457 | 412.91 | NA |
| 4 | RNS-present | No | -211.070 | 438.655 | 19.885 |
| 4 | RNS-present | Yes | -210.970 | 450.555 | 31.785 |

| | | | | | |
|---|---|---|---|---|---|
| 5 | RNS-present | No | -212.033 | 472.210 | 53.44 |
| 5 | RNS-present | Yes | -213.012 | 481.112 | 62.34 |

**C.** Results of model fit tests for alternative topology

| Number of Rate Categories | "Precursor" state? | AICc | deltaAIC |
|---|---|---|---|
| 1 | NA | 465.996 | 51.615 |
| 2 | No | 418.746 | 4.365 |
| 2 | Yes | 416.748 | 2.367 |
| 3 | No | 416.038 | 1.657 |
| 3 | Yes | 414.381 | 0 |
| 4 | No | 424.814 | 10.433 |
| 4 | Yes | 424.437 | 10.056 |
| 5 | No | 434.914 | 20.553 |
| 5 | Yes | 428.137 | 13.756 |

**Extended Data Table 2.** List of hypothesized origins of RNS in the NFC. Clades that also include non-nodulating taxa due to loss of RNS are marked with *. Incongruences with previously published phylogenies are marked †. Comparison groups that are sister taxa with an ancestral absence of RNS are marked with ‡ except in those cases where they are taxa with secondary loss and their closest RNS-present relatives. §*Vouacapoua* is generally regarded as non-nodulating, but see Moreira (1992).

| Order | Family or Subfamily (Fabales) | Nodulating clade | Comparison | SI annotation |
|---|---|---|---|---|
| Fabales | Papilionoideae | meso-Papilionoideae (50-kb inversion clade)* | non-*Nissolia* Adesmia clade (+) and *Nissolia*(-)‡ | Sup.Data.3; 1 |
| Fabales | Papilionoideae | Atelioids and Swartzioids *sensu stricto** | Atelioids (+) and *Bocoa, Trischidium* (-)‡ | Sup.Data.3; 2 |
| Fabales | Papilionoideae | *Dussia* | Amburaneae exclusive of *Dussia* | Sup.Data.3; 3 |

| | | | | |
|---|---|---|---|---|
| Fabales | Caesalpinioideae | Mimosoid Clade + Tachigali Clade + Dimorphandra Clade + Peltophorum group + *Moldenhawera*[*] | *Anadenanthera*(+) and *Parkia* (-)[‡]; *Viguieranthus* (+?) and *Zapoteca* (-)[‡] | Sup.Data.4; 4 |
| Fabales | Caesalpinioideae | *Chamaecrista* | *Cassia/Senna* (Cassieae Clade) or *Vouacapoua*[‡,§] | Sup.Data.4; 5 |
| Fabales | Caesalpinioideae | *Melanoxylum* + *Recordoxylon* | *Cassia/Senna* (Cassieae Clade) or *Vouacapoua*[‡,§] | Sup.Data.4; 6 |
| Rosales | Rosaceae | Subfamily Dryadoideae | Rosaceae exclusive of subfamily Dryadoideae | Sup.Data.5; 7 |
| Rosales | Cannabaceae | *Parasponia* | *Trema* (paraphyletic) | Sup.Data.5; 8 |
| Rosales | Rhamnaceae | *Ceanothus* | *Colubrina* (paraphyletic)[†] | Sup.Data.5;9 |
| Rosales | Rhamnaceae | Tribe Colletieae | *Ziziphus* (paraphyletic)[†] | Sup.Data.5;10 |
| Rosales | Elaeagnaceae | Elaeagnaceae | Dirachmaceae | Sup.Data.5;11 |
| Fagales | Betulaceae | *Alnus* | Subfamily Coryloideae[†] | Sup.Data.6;12 |
| Fagales | Casuarinaceae | Casuarinaceae | Ticodendraceae + Betulaceae | Sup.Data.6;13 |
| Fagales | Myricaceae | Myricaceae exclusive of *Canacomyrica* | *Canacomyrica* | Sup.Data.6;14 |
| Cucurbitales | Datiscaceae | *Datisca* (monogeneric family) | Tetramelaceae | Sup.Data.7;15 |
| Cucurbitales | Coriariaceae | *Coriaria* (monogeneric family) | Corynocarpaceae | Sup.Data.7;16 |

**Extended Data Table 3.** List of hypothesized losses of RNS in the NFC. The comparison is "none" when the loss clade is sister to a species-rich clade that includes independent gain and loss events. *See Supplementary Note 3.

| Order | Family or Subfamily (Fabales) | Loss clade | Comparison | SI annotation |
|---|---|---|---|---|
| Fabales | Papilionoideae | *Bocoa,Trischidium* | Atelioids | Sup.Data.3; 17 |
| Fabales | Papilionoideae | *Nissolia* | Adesmia clade exclusive of *Nissolia* | Sup.Data.3; 18 |
| Fabales | Caesalpinioideae | *Zapoteca* | *Viguieranthus* | Sup.Data.4; 19 |
| Fabales | Caesalpinioideae | *Vouacapoua* | *Melanoxylum* + *Recordoxylon* | Sup.Data.4; 20 |
| Fabales | Caesalpinioideae | *Parkia* | *Anadenanthera* | Sup.Data.4; 21 |
| Fabales | Caesalpinioideae | *Newtonia* | The most inclusive clade that includes *Prosopis* and *Inga* but not *Newtonia* | Sup.Data.4; 22 |
| Fabales | Caesalpinioideae | *Adenanthera,Tetrapleura* | RNS+ *Adenanthera* group | Sup.Data.4; 23 |
| Fabales | Caesalpinioideae | *Mora* | *Dimorphandra** | Sup.Data.4; 24 |
| Fabales | Caesalpinioideae | *Arapatiella* | *Jacqueshuberia* | Sup.Data.4; 25 |
| Fabales | Caesalpinioideae | *Peltophorum* group | The most inclusive clade that includes *Diptychandra* and *Inga* but not *Peltophorum* | Sup.Data.4; 26 |

**Extended Data Table 4.** Confidence intervals for transition rates estimated under the best model for the main topology (Fig. 1). Note that rows 6-11 are identical to rows 12-17 because under an ARD HRM model as implemented in corHMM, transitions between rate categories must be equal across observed states.

| Transition | Description | Rate Category | ML rate | Confidence Interval |
|---|---|---|---|---|
| **(2,R1) -> (1,R1)** | RNS-Loss | Intermediary hidden state | 0.0000 | (0, 4.290e-09) |
| **(1,R1) -> (2,R1)** | RNS-Gain | Intermediary hidden state | 0.0977 | (0.032,0.180) |
| **(2,R2) -> (1,R2)** | RNS-Loss | Precursor | 1.6373 | (0.718,6.846) |
| **(1,R2) -> (2,R2)** | RNS-Gain | Precursor | 0.0000 | (0,9.942e-09) |
| **(2,R3) -> (1,R3)** | RNS-Loss | Non-precursor (absorbing) | 0.0006 | (0.00047,0.001) |
| **(1,R2) -> (1,R1)** | Gain of intermediary hidden state | NA | 0.0086 | (0.007,0.020) |
| **(1,R3) -> (1,R1)** | None | NA | 0.0000 | (0,6.928e-09) |
| **(1,R1) -> (1,R2)** | Loss of intermediary hidden state to precursor | NA | 0.0180 | (0.014, 0.042) |
| **(1,R3) -> (1,R2)** | None | NA | 0.0000 | (0,2.674734e-09) |
| **(1,R1) -> (1,R3)** | Loss of intermediary hidden state to absorbing non-precursor state | NA | 0.1116 | (0.078,0.200) |
| **(1,R2) -> (1,R3)** | Loss of precursor to absorbing non-precursor state | NA | 0.0271 | (0.017,0.037) |
| **(2,R2) -> (2,R1)** | None | NA | 0.0086 | (0.007,0.020) |

| | | | | |
|---|---|---|---|---|
| **(2,R3) -> (2,R1)** | None | NA | 0.0000 | (0,6.928e-09) |
| **(2,R1) -> (2,R2)** | None | NA | 0.0180 | (0.014, 0.042) |
| **(2,R3) -> (2,R2)** | None | NA | 0.0000 | (0,2.674734e-09) |
| **(2,R1) -> (2,R3)** | None | NA | 0.1116 | (0.078,0.200) |
| **(2,R2) -> (2,R3)** | None | NA | 0.0271 | (0.017,0.037) |

**Extended Data Table 5.** Summary of taxonomic sampling of the NFC. Number of genera and species assignment to genera is based on The Plant List (version 1.1; http://www.theplantlist.org) except for Leguminosae subfamilies, which follow LPWG (2017).

| **Order** | **Family or Leguminosae Subfamily (Fabales)** | **Number of samples** | **Genera sampled/Total genera** |
|---|---|---|---|
| Fabales | Papilionoideae | 5121 | 443/503 |
| Fabales | Caesalpinioideae | 2100 | 146/148 |
| Fabales | Dialioideae | 38 | 16/17 |
| Fabales | Detarioideae | 367 | 72/84 |
| Fabales | Cercidoideae | 161 | 9/12 |
| Fabales | Duparquetioideae | 1 | 1/1 |
| Fabales | Polygalaceae | 320 | 22/27 |
| Fabales | Surianaceae | 3 | 4/4 |
| Fabales | Quillajaceae | 2 | 1/1 |
| Rosales | Rosaceae | 1917 | 99/109 |
| Rosales | Cannabaceae | 70 | 9/9 |
| Rosales | Rhamnaceae | 558 | 51/57 |
| Rosales | Moraceae | 698 | 44/49 |
| Rosales | Elaeagnaceae | 63 | 3/4 |
| Rosales | Ulmaceae | 41 | 7/8 |

| | | | |
|---|---|---|---|
| Rosales | Urticaceae | 642 | 50/58 |
| Rosales | Dirachmaceae | 1 | 1/1 |
| Rosales | Barbeyaceae | 1 | 1/1 |
| Fagales | Betulaceae | 92 | 6/6 |
| Fagales | Casuarinaceae | 37 | 4/4 |
| Fagales | Nothofagaceae | 12 | 1/1 |
| Fagales | Juglandaceae | 75 | 11/13 |
| Fagales | Ticodendraceae | 2 | 1/1 |
| Fagales | Fagaceae | 417 | 7/9 |
| Fagales | Myricaceae | 12 | 4/5 |
| Cucurbitales | Datiscaceae | 2 | 1/1 |
| Cucurbitales | Apodanthaceae | 3 | 3/3 |
| Cucurbitales | Anisophylleaceae | 5 | 2/4 |
| Cucurbitales | Corynocarpaceae | 2 | 1/1 |
| Cucurbitales | Coriariaceae | 5 | 1/1 |
| Cucurbitales | Cucurbitaceae | 343 | 95/134 |
| Cucurbitales | Tetramelaceae | 1 | 2/2 |
| Cucurbitales | Begoniaceae | 197 | 2/2 |

A

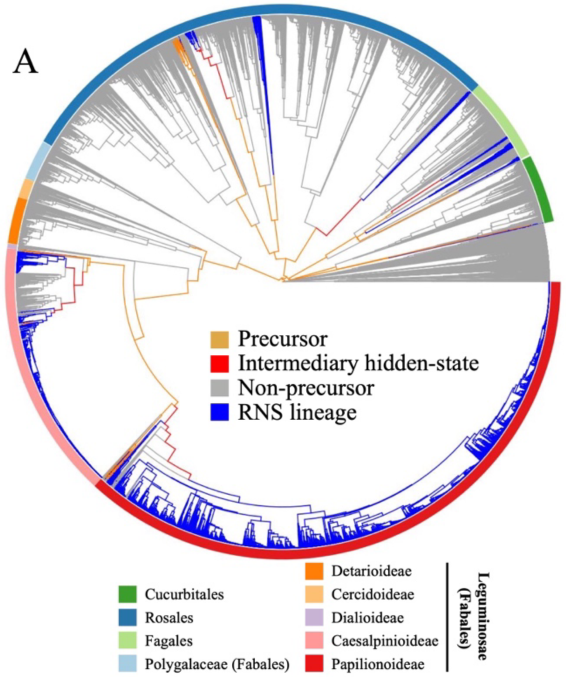


B

| | (1,R1) | (2,R1) | (1,R2) | (2,R2) | (1,R3) | (2,R3) |
|---|---|---|---|---|---|---|
| (1,R1) | - | .0977 | .0180 | NA | .1116 | NA |
| (2,R1) | .0000 | - | NA | .018 | NA | .1116 |
| (1,R2) | .0086 | NA | - | .000 | .0271 | NA |
| (2,R2) | NA | .0086 | 1.6373 | - | NA | .0271 |
| (1,R3) | .0000 | NA | .0000 | NA | - | NA |
| (2,R3) | NA | .0000 | NA | .000 | .0006 | - |

**Extended Data Figure 1.** Results of transition rate estimation and ancestral character state reconstruction based on an alternative NFC backbone topology (see Supplementary Note 4). A. Character states inferred by joint estimation of ancestral character states. Non-NFC clades are scaled down (unlabeled section) to highlight the NFC. NFC orders and legume subfamilies are indicated by colored bars. Branches are colored by the state estimated at their tipward node. B. Transition rate matrix of estimated rates from states listed on left to states listed at top. Rows are colored to correspond to colors in panel A.


A

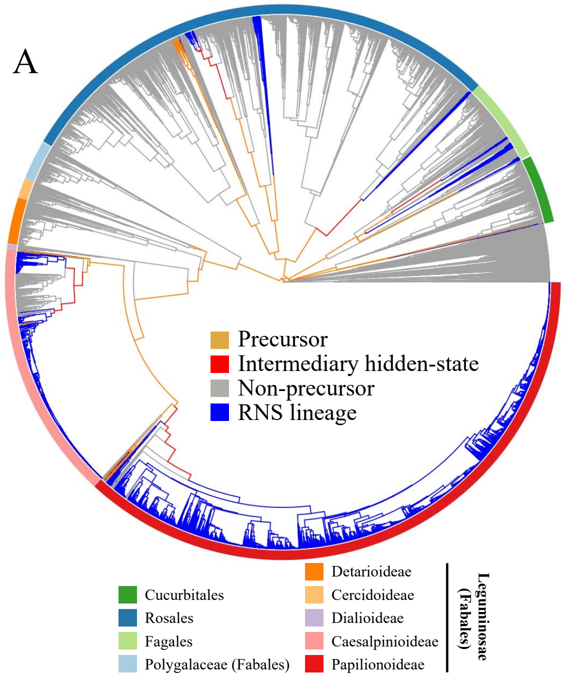


B

| | (1,R1) | (2,R1) | (1,R2) | (2,R2) | (1,R3) | (2,R3) |
|---|---|---|---|---|---|---|
| (1,R1) | - | .0977 | .0180 | NA | .1116 | NA |
| (2,R1) | .0000 | - | NA | .018 | NA | .1116 |
| (1,R2) | .0086 | NA | - | .000 | .0271 | NA |
| (2,R2) | NA | .0086 | 1.6373 | - | NA | .0271 |
| (1,R3) | .0000 | NA | .0000 | NA | - | NA |
| (2,R3) | NA | .0000 | NA | .000 | .0006 | - |

**Extended Data Figure 2.** Results of transition rate estimation and ancestral character state reconstruction based on the two-rate precursor model (deltaAIC 1.063). A. Character states inferred by joint estimation of ancestral character states. Non-NFC clades are scaled down (unlabeled section) to highlight the NFC. NFC orders and legume subfamilies are indicated by colored bars. Branches are colored by the state estimated at their tipward node. B. Transition rate matrix of estimated rates from states listed on left to states listed at top. Rows are colored to correspond to colors in panel A.

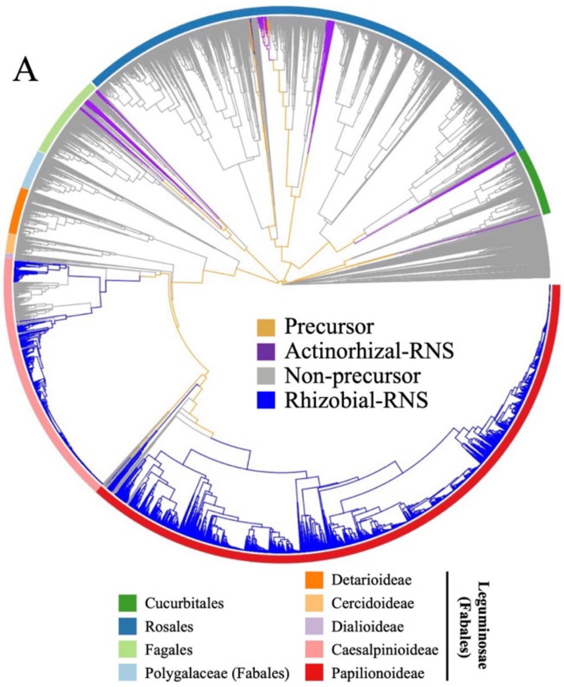


B

| | (1,R1) | (2,R1) | (3,R1) | (1,R2) | (2,R2) | (3,R2) |
|---|---|---|---|---|---|---|
| (1,R1) | - | .0045 | NA | .0374 | NA | NA |
| (2,R1) | .0643 | - | .0235 | NA | .0374 | NA |
| (3,R1) | .0134 | NA | - | NA | NA | .0374 |
| (1,R2) | .0000 | NA | NA | - | NA | NA |
| (2,R2) | NA | .0000 | NA | .0001 | - | .0000 |
| (3,R2) | NA | NA | .0000 | .0007 | NA | - |

**Extended Data Figure 3.** Results of transition rate estimation and ancestral character state reconstruction based on a model of ordered RNS-state change with three states (RNS-absent, Actinorhizal-RNS, and Rhizobial-RNS) and two rate categories (delta AIC 49.176). A. Character states inferred by joint estimation of ancestral character states. Non-NFC clades are scaled down (unlabeled section) to highlight the NFC. NFC orders and legume subfamilies are indicated by colored bars. Branches are colored by the state estimated at their tipward node. B. Transition rate matrix of estimated rates from states listed on left to states listed at top. Rows are colored to correspond to colors in panel A.

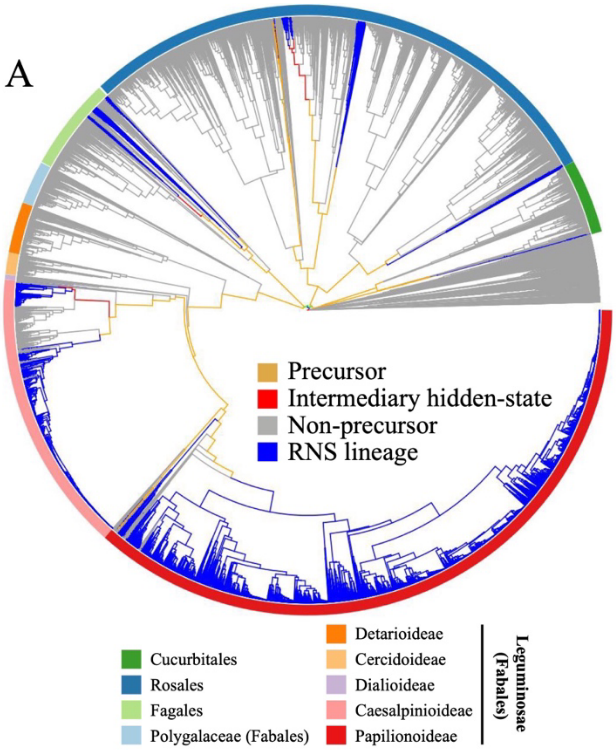


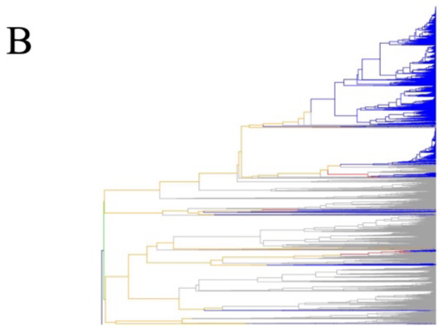


C

| | (1,R1) | (2,R1) | (1,R2) | (2,R2) | (1,R3) | (2,R3) |
|---|---|---|---|---|---|---|
| (1,R1) | - | .000 | .000 | NA | .000 | NA |
| (2,R1) | .001 | - | NA | .000 | NA | .000 |
| (1,R2) | .109 | NA | - | .100 | .022 | NA |
| (2,R2) | NA | .109 | .000 | - | NA | .022 |
| (1,R3) | .028 | NA | .008 | NA | - | NA |
| (2,R3) | NA | .028 | NA | .008 | .797 | - |

**Extended Data Figure 4.** Results of transition rate estimation and ancestral character state reconstruction in fixed-ancestral-state (RNS-present) analysis. A. Character states inferred by joint estimation of ancestral character states. Non-NFC clades are scaled down (unlabeled section) to highlight the NFC. NFC orders and legume subfamilies are indicated by colored bars. Branches are colored by the state estimated at their child node. B. A linear representation of the NFC phylogeny to highlight transitions in basal nodes that are not readily visible in the circular format; the taxon sequence is identical to that in the circle tree. C. Transition rate matrix of estimated rates from states listed on left to states listed at top. Rows are colored to correspond to colors in panel A.

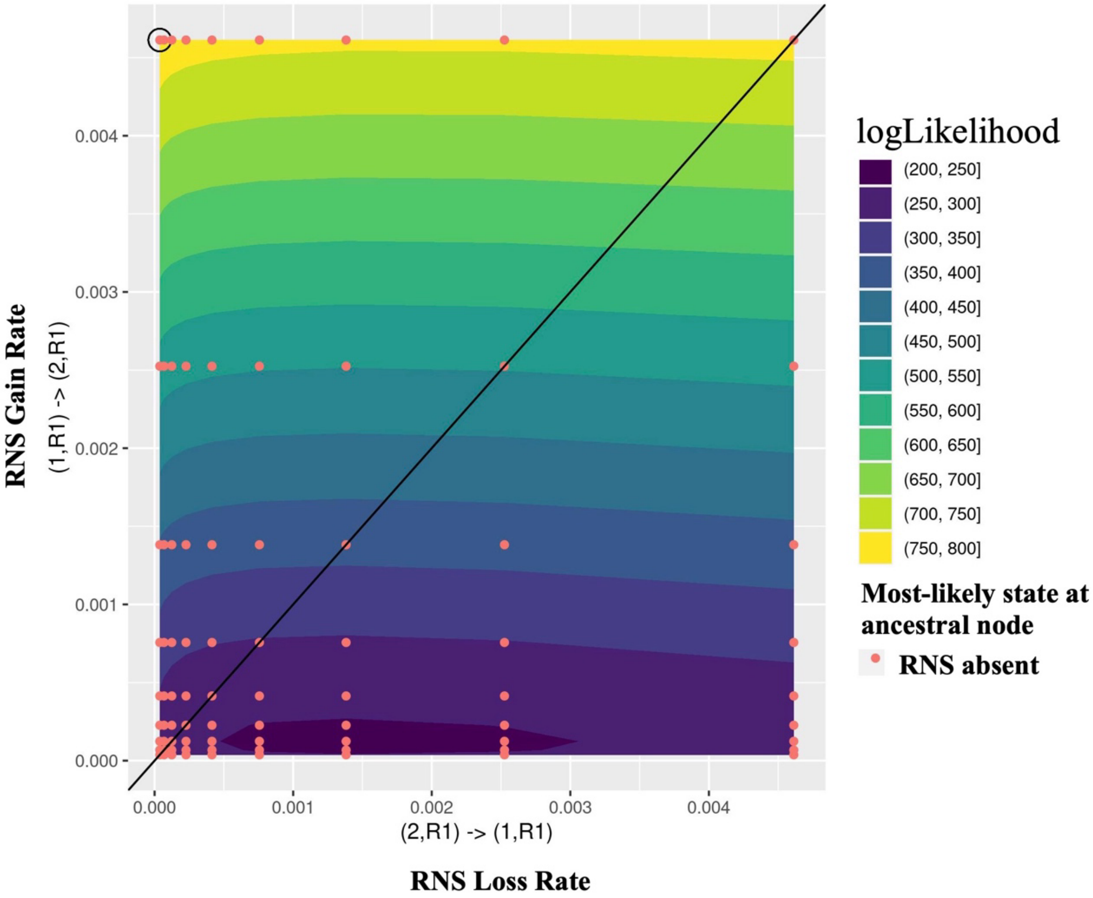


**Extended Data Figure 5.** Results of a set of simulated transition rates for a two-rate model of RNS gain and loss. Gain rate is on the X-axis, and loss rate is on the Y-axis. Plot background is colored by logLikihood scores for the plotted resulting model likelihoods; worse likelihoods are at the bottom, and better likelihoods are at the top. For each model, if the resulting marginal reconstruction estimated the NFC ancestor as RNS-absent, the plot point is colored red. (Ancestral RNS-presence would result in blue plot points, but no models yielded this result.)
